# *Arabidopsis thaliana* zinc accumulation in leaf trichomes is correlated with zinc concentration in leaves

**DOI:** 10.1101/2020.09.10.291880

**Authors:** Felipe K. Ricachenevsky, Tracy Punshon, David E. Salt, Janette P. Fett, Mary Lou Guerinot

## Abstract

Zinc (Zn) is a key micronutrient. In humans, Zn deficiency is a common nutritional disorder, and most people acquire dietary Zn from eating plants. In plants, Zn deficiency can decrease plant growth and yield. Understanding Zn homeostasis in plants can improve agriculture and human health. While root Zn transporters in plat model species have been characterized in detail, comparatively little is known about shoot processes controlling Zn concentrations and spatial distribution. Previous work showed that Zn hyperaccumulator species such as *Arabidopsis halleri* accumulate Zn and other metals in leaf trichomes. The model species *Arabidopsis thaliana* is a non-accumulating plant, and to date there is no systematic study regarding Zn accumulation in *A. thaliana* trichomes. Here, we used Synchrotron X-Ray Fluorescence mapping to show that Zn accumulates at the base of trichomes of *A. thaliana*, as had seen previously for hyperaccumulators. Using transgenic and natural accessions of *A. thaliana* that vary in bulk leaf Zn concentration, we demonstrated that higher leaf Zn increases total Zn found at the base of trichome cells. Furthermore, our data suggests that Zn accumulates in the trichome apoplast, likely associated with the cell wall. Our data indicates that Zn accumulation in trichomes is a function of the Zn status of the plant, and provides the basis for future studies on a genetically tractable plant species aiming at understanding the molecular steps involved in Zn spatial distribution in leaves.

## Introduction

Zinc (Zn) is an essential micronutrient, serving as a cofactor for many enzymes and transcription factors (Maret, 2009). An estimated 8% of the *Arabidopsis thaliana* proteome binds to Zn (Andreini et al., 2006). In humans, Zn deficiency is the second most widespread nutritional deficiency (after iron), with estimated 25% of the population at risk of low Zn intake, especially when the diet is composed mostly of cereal grains (Maret and Sandstead, 2006;Gomez-Galera et al., 2010). Since plants are the primary source of Zn entry into the food chain, understanding Zn homeostasis is essential to produce plants that accumulate more Zn (and other nutrients) in their edible tissues (Ricachenevsky et al., 2015;Garcia-Oliveira et al., 2018).

The genetic regulation of Zn uptake, distribution, detoxification and storage in *A. thaliana* has been functionally characterized in some detail (Sinclair and Kramer, 2012;Ricachenevsky et al., 2015). Most of our knowledge is focused on root control of Zn acquisition and Zn excess detoxification. Primary Zn uptake are from the soil is likely performed by genes of the ZIP (<u>Z</u>inc-regulated/<u>I</u>ron-regulated transporter <u>P</u>rotein) family (Assuncao et al., 2010;Milner et al., 2013;Sinclair et al., 2018). In rice, OsZIP9 was as recently described as responsible for Zn acquisition from the rhizosphere (Huang et al., 2020;Tan et al., 2020;Yang et al., 2020). The ZIP transporters were the first Iron (Fe) and Zn transporters characterized in plants, and were shown to transport Fe, Zn and other divalent cations in several plant species (Eide et al., 1996;Tiong et al., 2014;Evens et al., 2017;Ricachenevsky et al., 2018). The *A. thaliana* bZIP19 and bZIP23 transcription factors control the Zn deficiency response, in which ZIP transporters are upregulated (Assuncao et al., 2010;Inaba et al., 2015;Lilay et al., 2018). AtZIP4 and AtZIP9, two direct targets of bZIP19 and bZIP23, are up regulated in roots in response to a local Zn deficiency signal. On the other hand, AtMTP2 (<u>M</u>etal <u>T</u>olerance <u>P</u>rotein 2), an endoplasmic reticulum (ER)-localized Zn transporter, is also up regulated in roots, but in response to a shoot-derived Zn deficiency signal (Sinclair et al., 2018). Other members of this family, vacuolar transporters such as AtMTP1 (Kobae et al., 2004;Desbrosses-Fonrouge et al., 2005) and AtMTP3, detoxify excess Zn by sequestering it into the vacuole. ZIF-Like (<u>Z</u>inc-<u>I</u>nduced <u>F</u>acilitator) family transporters may be involved Zn tolerance, either through transport of Zn bound to nicotianamine (AtZIF1(Haydon et al., 2012)) or direct Zn sequestration (AtZIF2(Remy et al., 2014)) in vacuoles. Moreover, Zn efflux transporters such as AtHMA2 and AtHMA4 (<u>H</u>eavy <u>M</u>etal-<u>A</u>ssociated transporters) perform Zn loading from the root symplast into the xylem for long distance transport (Hussain et al., 2004), and are also involved in Zn loading in seeds (Olsen et al., 2016).

Zn accumulation and storage in shoots, on the other hand, is not well understood. Previous work in Zn hyperaccumulator species *Arabidopsis halleri* and *Nocceae caerulescens* (Gonneau et al., 2017;Stein et al., 2017;Schvartzman et al., 2018) which are close relatives of *A. thaliana*, show how critical Zn homeostasis genes are in establishing their remarkable tolerance to Zn levels that are lethal to other species. Both *A. halleri* and *N. caerulescens* rely on multiple copies of genes *MTP1* and *HMA4* for their Zn hyperaccumulation phenotypes (Hanikenne et al., 2008;Shahzad et al., 2010;S et al., 2011). Evidence indicates that Zn accumulation might be related to herbivory deterrence (Kazemi-Dinan et al., 2014;Martos et al., 2016). Interestingly, *A. halleri* accumulates high concentration of Zn in its trichomes. This accumulation occurs specifically at the base of tricomes as a narrow ring (Kupper et al., 2000; Zhao FJ, 2000; Sarret et al., 2009; Sarret et al., 2009) Elements such as cadmium (Cd), for which *A. halleri* is also hypertolerant, also accumulate in trichomes (Isaure et al., 2015). Because the hyperaccumulator/hypertolerant species *A. halleri* and the non-hyperaccumulator *A. lyrata* both accumulate Zn and Cd in trichomes (Sarret et al., 2009;Isaure et al., 2015) in a similar pattern, it seems unlikely that this is a hyperaccumulation mechanism.

*A. thaliana* non-glandular tricomes are derived from epidermal cells that undergo endoreduplication, which consists of replication of the genome without mitosis, and results in increased cell size (Uhrig and Hulskamp, 2010). These trichomes differ from the secretory trichomes, such as those found in tobacco, which were shown to secrete Cd and Zn (Sarret et al., 2006; Isaure MP, 2010). Despite not having a secretory function, non-glandular trichomes are likely hotspots for metal accumulation (Blamey et al., 2015;Kopittke et al., 2018). Cd has been detected in *A. thaliana* non-glandular trichomes (Ager et al., 2003; Huguet et al., 2012). Sunflower (*Helianthus annuus*) non-glandular trichomes accumulate Mn when the metal is in excess in the growth medium (Blamey et al., 2015), while Zn accumulates rapidly at the trichome base when Zn is sprayed on leaves (Li et al., 2019), a phenomenon also observed in soybean (Li et al., 2018). Despite the work on *A. lyrata* and *A. halleri*, little is known about metal accumulation in the trichomes of the model species *A. thaliana*.

Our aim in this study was to understand how Zn availability affects *A. thaliana* Zn accumulation in trichomes and in leaves. To do this we conducted elemental mapping using synchrotron X-ray fluorescence (SXRF) of both *A. thaliana* natural variants and transgenic plants provided with a variety of Zn concentrations in the growth medium. Non-glandular trichomes accumulated more Zn as leaf Zn concentrations increased and Zn accumulation at the base of the trichome was observed, likely localized in the trichome cell apoplast. Our results suggest that plants may actively change the partitioning of Zn to trichomes in response to leaf Zn supply.

## Materials and Methods

### Plant materials and growth conditions

For growth in axenic conditions, seeds were sterilized for 15 minutes in 25% NaOH and 0.05% SDS, washed 5 times in sterile H_2_O and stratified at 4°C for three days. Sterile 0.1% agar was used to suspend seeds, which were sown using a pipette onto plates made with full strength Gamborg’s B5 media plus vitamins, 1mM MES (2-(N-morpholino)ethanesulfonic acid), 2% sucrose and 0.6% agar. After five days, seedlings were transferred to minimal media containing 2 mM MES, 2 mM Ca(NO_3_)_2_.4H_2_O, 0.75 mM K_2_SO_4_, 0.65 mM MgSO_4_.7H_2_O, 0.1 mM KH_2_PO_4_, 10 µM H_3_BO_3_, 0,1 µM MnSO_4_, 50 nM CuSO_4_, 5 nM (NH_4_)_6_Mo_7_O_24_ and 50 µM Fe-EDTA. ZnSO_4_ was added to a final concentration of 50 nM in control conditions, or at indicated concentrations (50 µM or 100 µM). Plates were kept at 22°C with 16 hours of light/ 8 hours of dark in growth chambers. Seedlings were analyzed after 15 days of growth.

### Preparation of samples and mapping by two-dimensional XRF

In all experiments, we compared the detached 7th true leaf of each plant cultivated under the conditions described in the methods. Lower resolution (7 µm steps) 2D elemental mapping was conducted on fresh, unfixed leaf samples placed between kapton films. Elemental maps were collected during multiple experiments at beamline 2-3 of the Stanford Synchrotron Radiation Lightsource (SSRL). Beam line 2-3 uses a water-cooled double crystal monochromator with either a Si(220) or Si(111) crystal and a Vortex single element detector. The beam was focused using a Pt-coated Kirkpatrick-Baez mirror pair (Xradia Inc.) and tuned to 11 keV. All maps were collected from the 7th true leaf. Elemental mapping was performed in 7 µm steps with a 50 ms dwell time for whole leaf maps, and in 2µm steps and 50 ms dwell time for trichome 2D mapping. The XRF maps were analyzed using the Microanalysis Toolkit (http://home.comcast.net/~sam_webb/smak.html). To determine total abundance of elements in whole leaf maps and in individual trichomes, region of interest (ROI) analysis was performed. Background counts in each trichome ROI were calculated by multiplying the mean counts per pixel from the whole leaf map by the number of pixels in ROI, and then subtracting this value from the total abundance. Percentage of elements in trichomes was calculated by the sum of total abundance in each trichome divided by total abundance in whole leaf map.

High-resolution elemental mapping (0.6 µm steps) of trichomes was conducted on embedded, sectioned trichomes from the 7th true, fully expanded leaf, using methods optimized for minimizing elemental distribution during fixation (Punshon et al., 2012). High-resolution X-ray fluorescence microscopy was performed at Beamline 2-ID-D of the Advanced Photon Source at the Argonne National Laboratory (Cai Z, 2000). An incident X-ray energy of 10 keV was chosen to excite elements from P to Zn. A Fresnel zone plate focused the X-ray beam to a spot size of 0.2 × 0.2 µm on the sample, which was raster scanned (Yun W, 1999) at resolutions of 0.6 µm step in the trichome plus mesophyll map, with dwell times ranging from 0.5 to 1 s per pixel. X-ray fluorescence from the sample was captured with an energy-dispersive silicon drift detector. Data is expressed as normalized fluorescence counts.

### Statistical analyses

When appropriate, data were subjected to ANOVA and means were compared by the Tukey HSD or Student’s t-test using the JMP 10.0 for Mac (SAS Inc., USA).

## Results

### OsZIP7 constitutive expression leads to altered Zn accumulation in leaves

We previously showed that OsZIP7 expression under the control of 35S promoter in *A. thaliana* leads to increased Zn concentration in leaves (Ricachenevsky et al., 2018). To investigate alterations in Zn localization, we used two-dimensional SXRF mapping of leaves of wild type (WT) and *OsZIP7*-expressing plants (hereafter OsZIP7-FOX) to identify possible changes in Zn distribution. This technique provides information about all Zn regardless of chemical speciation. Leaves of plants grown under control conditions showed Zn evenly distributed throughout the leaf, with higher concentrations in hydathodes and closer to the petiole detachment site (Figure 1A). In leaves of OsZIP7-FOX, Zn was found highly concentrated in small, punctuated areas at the leaf surface (Figure 1A). When Zn localization was overlaid with localization of Ca, which is known to accumulate in leaf trichome papillae (Esch et al., 2003), it was clear that Zn was accumulating at the base of trichomes in leaves of OsZIP7-FOX plants, a distribution that was not observed in WT (Figure 1B). Leaves from *glabra-1* mutant plants (Hulskamp et al., 1994), which lack trichomes, showed a uniform Zn distribution across the leaf surface similar to wild type when plants are grown with no added Zn (Figure 1A and 1B). However, when plants were grown on media supplemented with 50 µM or 100 µM Zn, the punctate pattern of Zn distribution associated with the base of trichomes was observed in both WT and OsZIP7-FOX leaves (Figure 1C-1F) wheras leaves of the trichome-less *glabra-1* mutant showed evenly distributed Zn and Ca. This confirms that Zn is accumulating at the base of trichomes, as observed for *A. lyrata* and *A. halleri* (Kupper et al., 2000;Zhao FJ, 2000).

**Figure 1.**
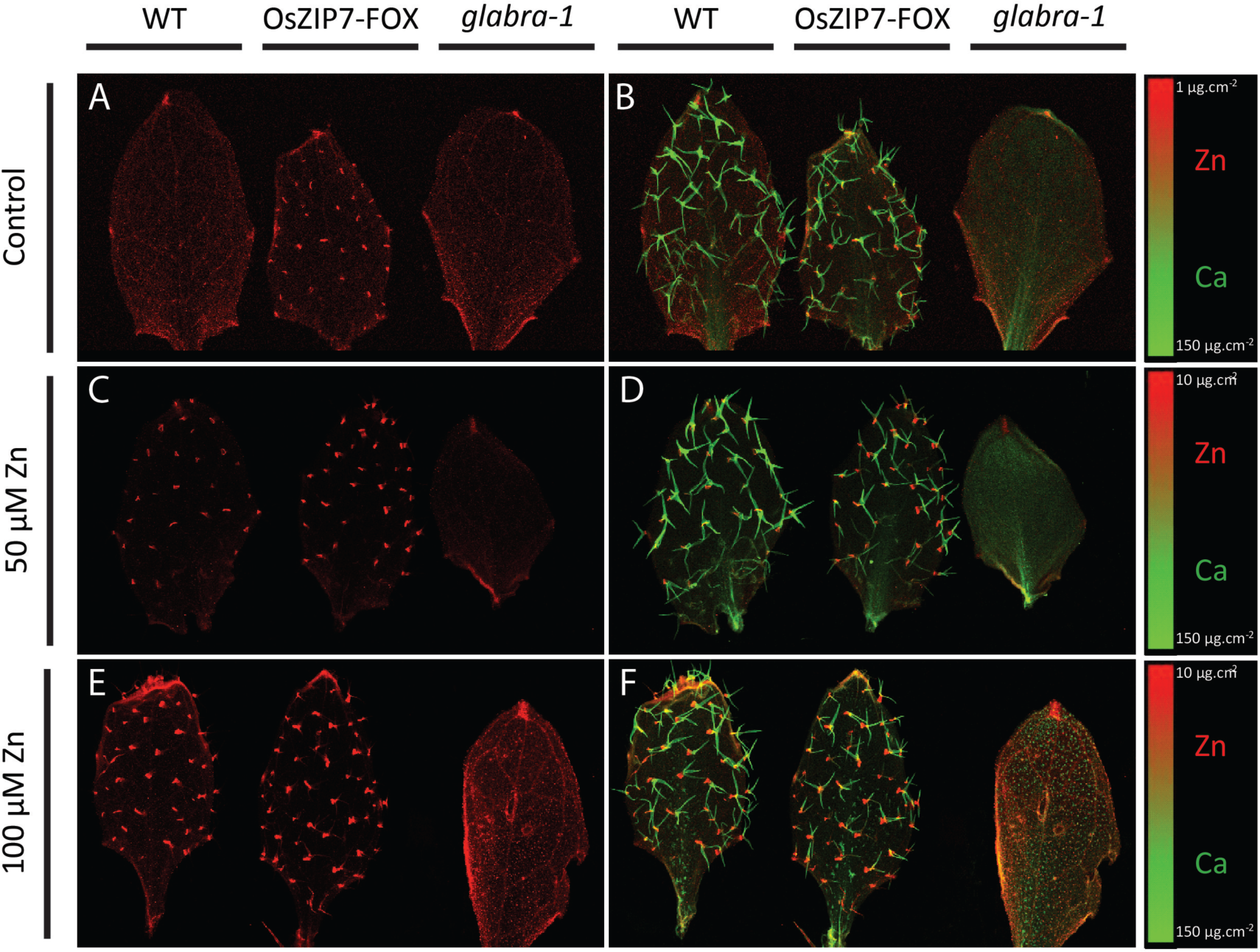
Elemental maps of Col-0, OsZIP7-OE and the *glabrous-1* mutant leaves. Maps show Zn localization (in red) in leaves of plants grown under control (A), 50 µM Zn (C) and 100 µM Zn (E), and Zn and Ca (in green) overlay in leaves from plants grown under control (B), 50 µM Zn (D) and 100 µM Zn (F).

Region of interest (ROI) analysis allowed us to dtermine total Zn per trichome from mapping data collected via SXRF. The OsZIP7-FOX leaf grown under control conditions had the lowest total Zn per trichome, considering the maps in which Zn in trichomes is observed (i.e., excludes WT Col-0 leaves under control conditions; Figure 2). In leaves of plants growing at 50 µM Zn, total Zn per trichome was higher in both WT and OsZIP7-FOX compared to OsZIP7-FOX under control conditions. The total Zn per trichome in WT and OsZIP7-FOX grown with 50 µM Zn were not significantly different (Figure 2). However, comparing WT and OsZIP7-FOX leaves from plants grown with 100 µM Zn, OsZIP7-FOX trichomes had clearly higher total Zn per trichome (Figure 2). We showed previously that OsZIP7-FOX lines accumulate higher Zn concentration in their leaves compared to WT under these conditions (Ricachenevsky et al., 2018). Our data indicate that OsZIP7 expression in *A. thaliana*, which leads to increased accumulation of Zn in leaves, also leads to accumulation of Zn at the base of trichomes.

**Figure 2.**
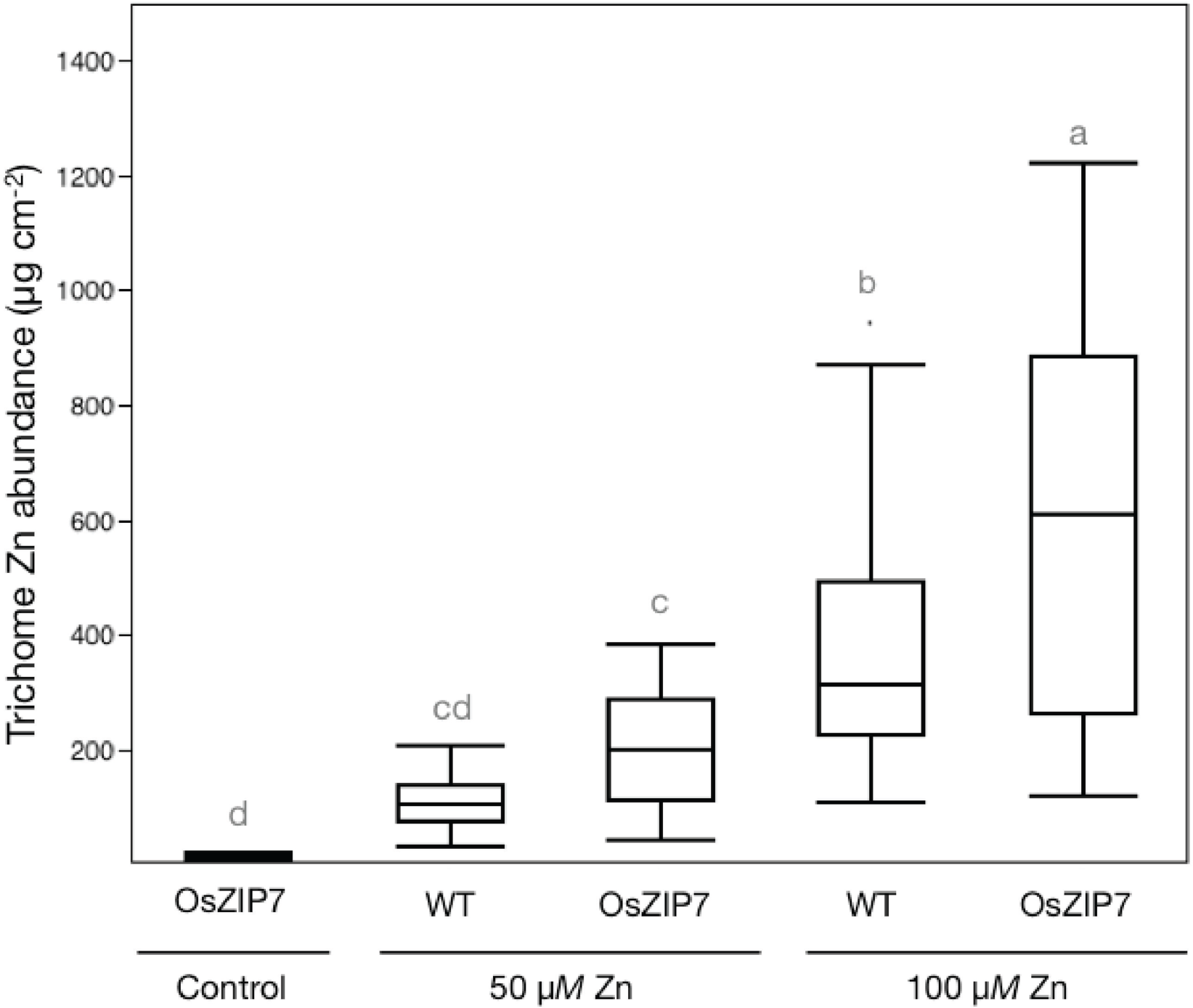
Total Zn abundance in individual trichomes. Significant differences detected by one-way ANOVA and Tukey HSD are represented by different lower case letters above the boxplots.

### Trichomes accumulate Zn in a ring around the base in *Arabidopsis thaliana*

To gain more information on the nature of the characteristic pattern of Zn accumulation at the base of trichomes, we used higher-resolution SXRF mapping of fresh trichomes. Plants grown at 50 µM Zn were chosen because both WT and OsZIP7-FOX leaves accumulate Zn in trichomes under these conditions (Figure 1C and 1D), and this concentration was sub-toxic to *A. thaliana* plants under our experimental conditions (Ricachenevsky et al., 2018). Zn localization maps of WT leaves showed a ring-shaped pattern around the base of the trichome (Figure 3A and 3B). In the mapped trichome of the OsZIP7-FOX plant, the Zn ring was thicker than in the mapped trichome of the WT plant, appearing to increase its domain farther away from the trichome base and up into the stalk (Figure 3C and 3D). Because these are individual trichomes, and the median total Zn per trichome from plants cultivated under these conditions are not statistically different (Figure 2), the distinct Zn distribution pattern observed is likely found in trichomes of both WT and OsZIP7 plants. Therefore, our data show that Zn accumulation at the base of the trichome occurs in a ring shape, which can vary in thickness depending on Zn concentration.

**Figure 3.**
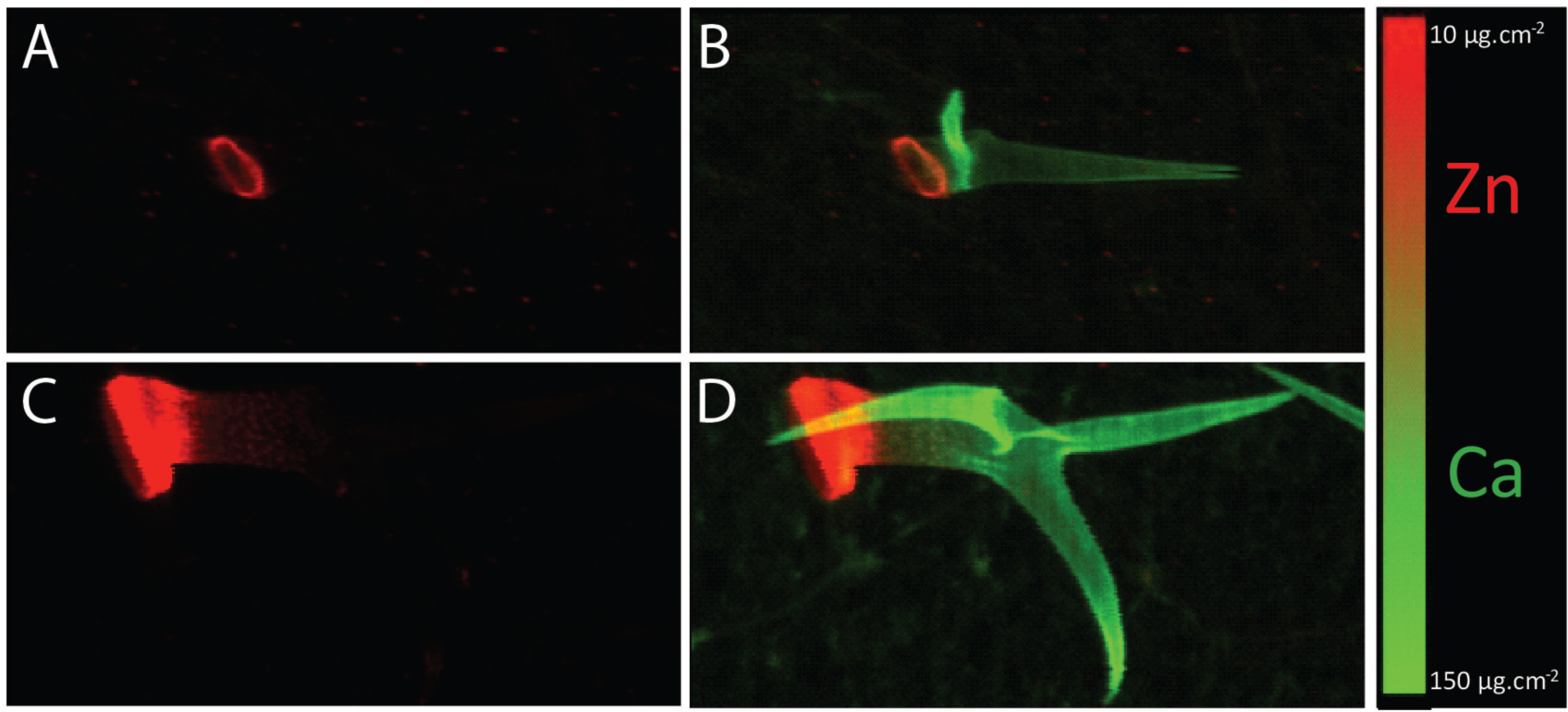
High-resolution elemental maps of individual fresh hydrated trichomes. Maps show Zn (in red) in trichomes of Col-0 (A) and OsZIP7-OE (C), and Zn and Ca (in green) localization overlay of Col-0 (B) and OsZIP7-OE (D).

### Zn localized in rings at the trichome base is likely to be extracellular

Next, we asked whether the ring-shaped Zn accumulation was at the base of the trichome cell or in the surrounding socket cells (Figure 4A, black arrows), which attach the trichome to the epidermis (Schliep et al., 2010). To answer that question, SXRF maps of longitudinally sectioned trichomes were collected from OsZIP7-FOX plants grown on control or 50 µM Zn (Figure 4A) since trichomes of this genotype have Zn in both conditions (Figure 2). In trichomes from control conditions, the trichome cell showed the highest Zn fluorescence, compared to socket cells or epidermal cells (Figure 4B). In trichomes from plants grown with 50 µM Zn, the trichome cell showed markedly higher Zn fluorescence than controls, while socket cells and epidermal cells had only marginal variations (Figure 4C-E), demonstrating that the trichome, not the underlying cells, is the primary site of Zn accumulation. We also observed stretches of lower Zn fluorescence inside the trichome cell (Figure 4E, white arrows), indicating possible Zn localization in the cytoplasm, since this Zn pattern is consistent with cytoplasm localization (Gutierrez-Alcala et al., 2000). Therefore, the regions with a high Zn fluorescence signal (Fig 4D, white arrows) are likely the apoplast. Moreover, we observed that chloroplasts are a major site of Zn localization in mesophyll cells (Figure 4D and 4E, yellow arrows).

**Figure 4.**
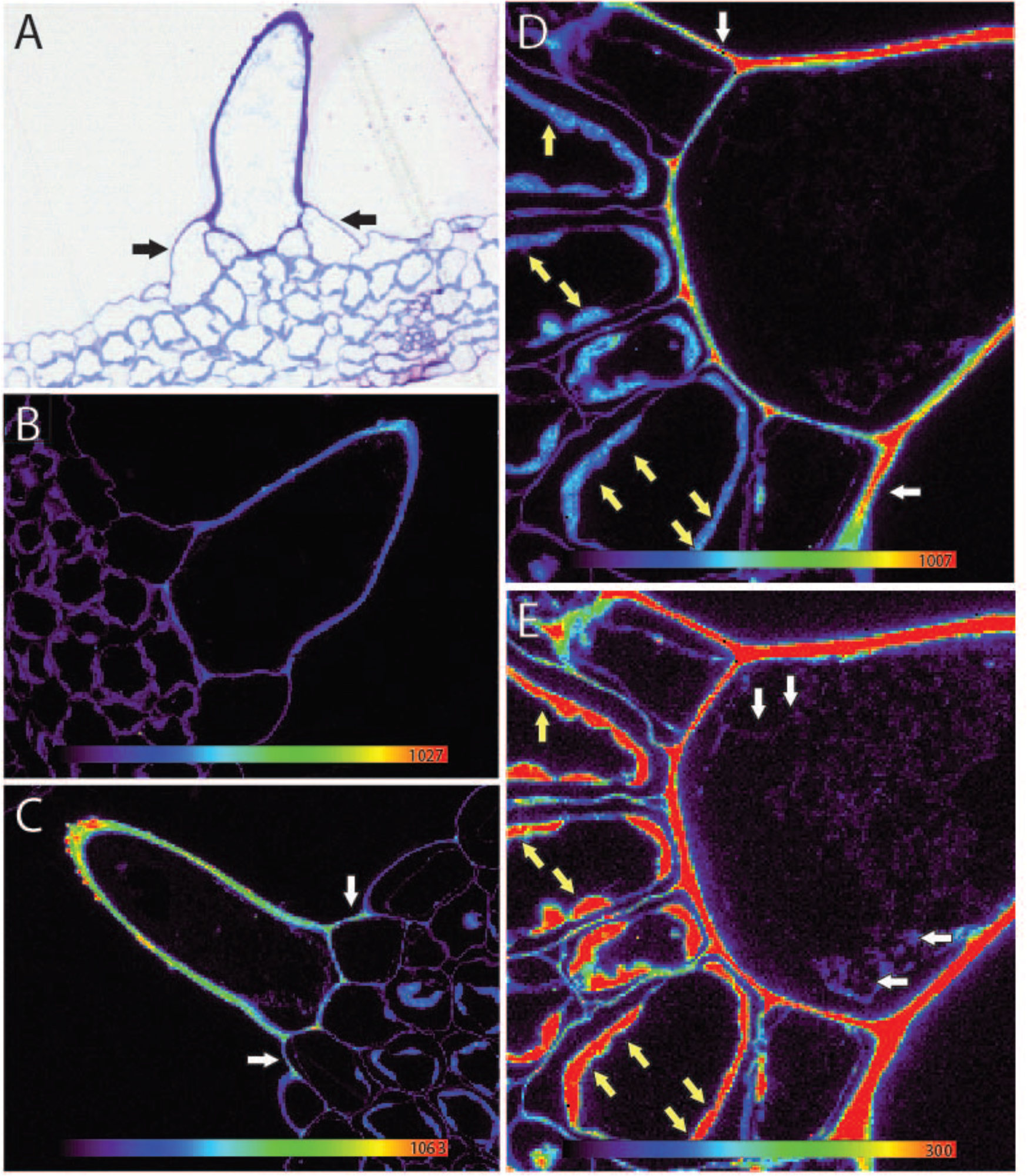
High-resolution elemental maps of trichomes sectioned in the sagittal plane. (A) Shows the sectioned trichomes used. Zn localization is shown in a trichome grown under control conditions (B) and under 50 µM Zn (C). (D) Zn localization in a trichome from a plant grown under 50 µM Zn scaled to the highest pixel abundance and (E) scaled to exclude the top two thirds of the data to allow low-abundance Zn distributions to be seen more clearly. White arrows in (C) and (D) show regions where Zn is in the apoplast of the trichome, suggesting that Zn moves from the trichome cell to surrounding cells. White arrows in (E) show features resembling cytoplasmic stretches within trichome cells, with lower Zn abundance. Yellow arrows in (E) indicate Zn localization in chloroplasts.

### Variations in leaf Zn concentration influence accumulation at the base of trichomes in *A. thaliana* natural accessions

The first part of this study showed that *OsZIP7* expression in *A. thaliana* led to increased Zn accumulation at the base of trichomes (Figure 1-3). We then asked whether the ability to accumulate different Zn concentrations in leaves would affect Zn in trichomes. To answer this question, we used Zn concentrations data from leaves of 349 *A. thaliana* natural accessions (www.ionomicshub.org; (Baxter et al., 2007)), and selected a subset of accessions with high and low Zn concentrations (Table 1). We mapped leaves of the accessions with the highest and the lowest Zn concentrations, Kn-0 and Fab-2, respectively (Figure 5, Table 1). Fab-2 was recently shown to harbor a loss of function allele for *AtHMA4*, which decreases Zn translocation from roots to leaves (Chen et al., 2018). Col-0 was included as the reference accession. Fab-2 leaves had several trichomes without Zn, and others with very low Zn fluorescence (Figure 5C and 5D). Conversely, Kn-0 showed higher Zn fluorescence in its trichomes compared to Col-0 (Figure 5A, 5B, 5E and 5F). ROI analyses of these maps showed that Kn-0 had higher total Zn per trichome, whereas Fab-2 trichomes showed lower total Zn per trichome, compared to the reference accession Col-0 (Figure 5G). These data suggest that increased levels of Zn in leaves are correlated with increase Zn accumulation at the base of the trichome.

**Table 1.** Natural accessions of *A. thaliana* used in this work. The ten highest and ten lowest Zn concentration in leaves are shown, from the 349 accessions found at the Ionomics Atlas. Accessions used are in bold and underlined.

| Group | Accession | Zn concentration (µg/g)<sup>a</sup> |
| --- | --- | --- |
| Low Zn accessions | **<u>Fab-2</u>** | 41.15 |
|  | **<u>Rev-2</u>** | 70.44 |
|  | **<u>Pn-0</u>** | 78.6 |
|  | **<u>Ste-3</u>** | 79.31 |
|  | TAD 01 | 79.51 |
|  | **<u>T1060</u>** | 80.03 |
|  | **<u>Shahdara</u>** | 84.3 |
|  | Mc-0 | 84.54 |
|  | Wag-3 | 85.13 |
|  | **<u>TOU-E-11</u>** | 87.79 |
| Reference accession | **<u>Col-0</u>** | 131.92 |
| High Zn accessions | **<u>ZdrI 2-25</u>** | 178.39 |
|  | **<u>Ull2-5</u>** | 181.23 |
|  | **<u>Ob-1</u>** | 182.69 |
|  | **<u>PHW-13</u>** | 196.65 |
|  | Ors-2 | 196.76 |
|  | **<u>NC-6</u>** | 201.48 |
|  | **<u>Uod-7</u>** | 205.2 |
|  | **<u>Na-1</u>** | 205.38 |
|  | Sav-0 | 238.5 |
|  | **<u>Kn-0</u>** | 254.53 |

<sup>a</sup> Data generated by ICP-MS and downloaded from the ionomics Atlas (http://www.ionomicshub.org/ionomicsatlas/).

**Figure 5.**
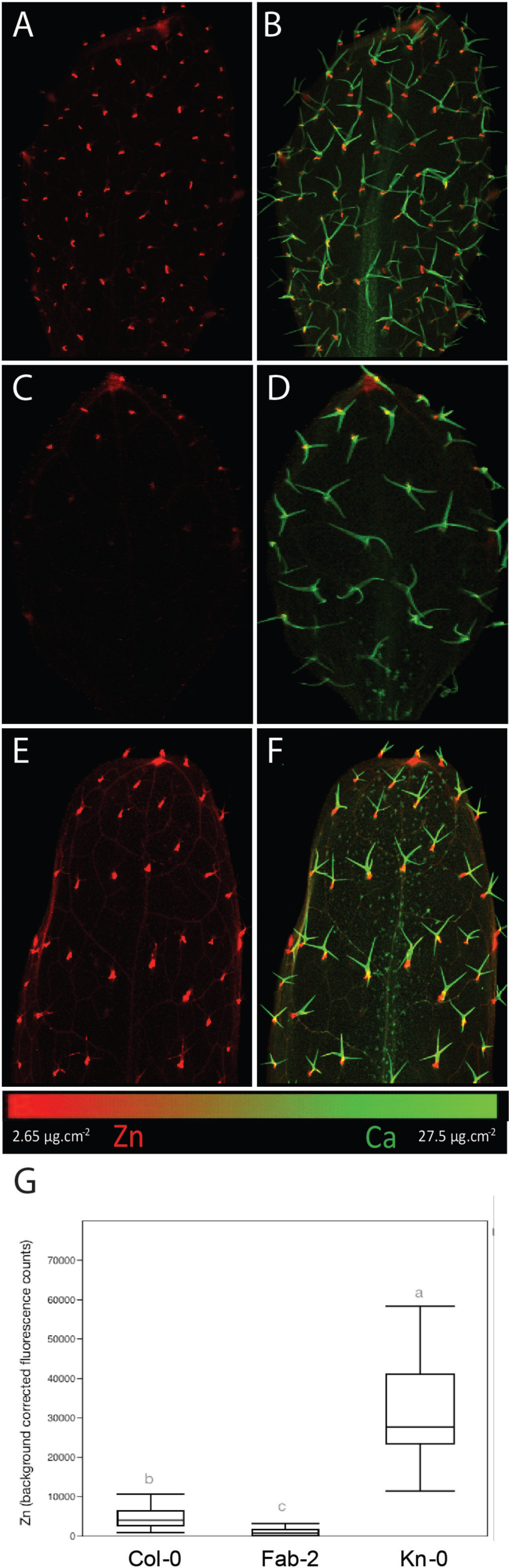
Total Zn per trichome of *A. thaliana* natural accessions with contrasting bulk leaf Zn concentrations. Total Zn abundance in trichomes was quantified using Region of Interest (ROI) analysis of leaves. Significant differences by one-way ANOVA and Tukey HSD are shown.

In addition to Col-0, Kn-0 and Fab-2, we mapped another 13 accessions with contrasting leaf Zn concentrations for a total of 16 accessions (Table 1). These accessions span the high and low ranges of Zn distribution in the iHUB, available at www.ionomicshub.org, allowing us to explore wide natural variation in leaf Zn concentration and its relationship to trichome Zn accumulation. Using ROI analyses, we determined the (1) mean count per pixel (a proxy for Zn concentration) per leaf; (2) percentage of total Zn sequestered within trichomes; and (3) percentage of Zn not associated with trichomes (Figure 6).

**Figure 6.**
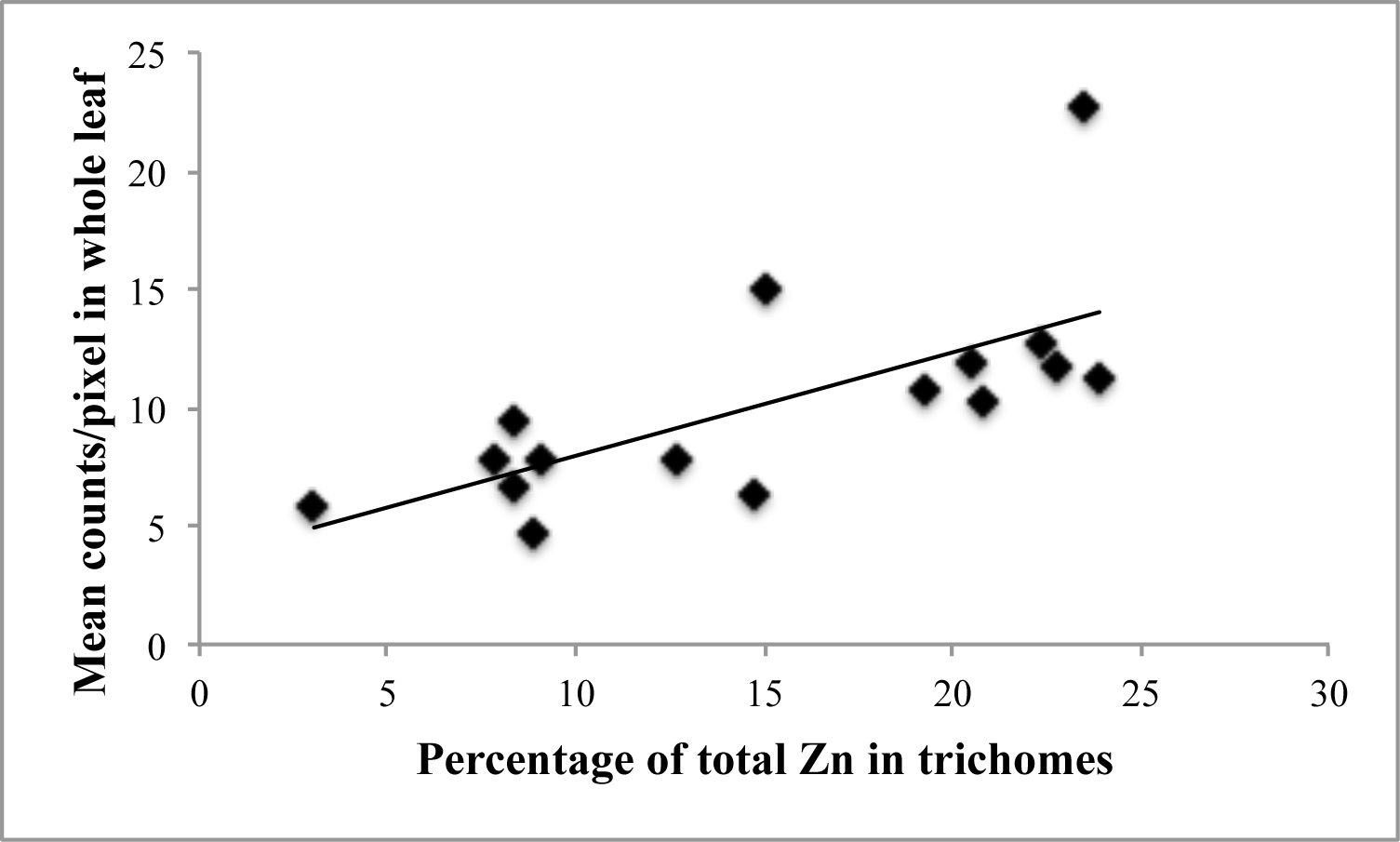
Zn accumulation in trichomes increases with Zn in leaves. Correlation of the percentage of the total Zn in trichomes with mean Zn counts in whole leaf of 16 *A. thaliana* accessions.

Next, we compared the mean fluorescence counts per pixel with the percentage of total Zn found in trichomes, for each accession (Figure 6). We found a positive correlation (R = 0.6871, p = 0.003) between the two variables, and consequently a negative, inverse correlation of Zn abundance not in trichomes (i.e., elsewhere in the leaf surface) and mean count per pixel. These data support the observations that *A. thaliana* accessions with higher leaf Zn concentrations accumulate a higher percentage of Zn at the base of trichomes, with a lower percentage elsewhere in the leaf. We therefore propose that Zn accumulation at the base of trichomes increases with increased Zn accumulation in the whole leaf, indicating trichomes may be an important Zn allocation site in leaves with high Zn concentration.

## Discussion

### Zn accumulates at the base of trichomes in *A. thaliana*

Here we have demonstrated that *A. thaliana* accumulates Zn at the base of the trichome. In several plant species, trichomes are common sites of excess metal accumulation, including Pb, Zn and Cd in *Nicotiana tabacum* (Sarret et al., 2006; Isaure MP, 2010), Cd in *Brassica juncea* (Salt et al., 1995), Mn in *Helianthus annuus* (Blamey et al., 2015) and Ni in the hyperaccumulator species *Alyssum lesbiacum* (Kupper et al., 2001). In the Zn hyperaccumulator/hypertolerant *A. halleri*, the base of trichomes accumulates the highest concentrations of Zn in the leaves (Kupper et al., 2000; Zhao FJ, 2000). Despite being regions of high accumulation, trichomes do not account for the majority of Zn in leaves: a comparison with non-hyperaccumulator *A. lyrata* showed that Zn in trichomes of *A. halleri* accounts for 10% of the total, while *A. lyrata* trichomes account for 20% of the total (Sarret et al., 2009). The data for *A. lyrata* agrees with our findings in *A. thaliana*, another non-hyperaccumulator, with Zn accumulation in trichomes ranging from 4% to 23% (Figure 6).

Using resin-embedded sections of trichomes, we found that Zn accumulates mostly at the base of the trichome cell and not in socket cells, which attach the trichome to the epidermis (Figure 4). The clear localization of Zn in trichome cells corroborates what we observed using fresh samples (Figure 1, Figure 5), suggesting that our high-resolution maps are of biological significance, although embedding might cause metal redistribution (Kopittke et al., 2018;van der Ent et al., 2018). Our method has already been used to show changes in calcium (Ca) distribution in seeds of *cax1, cax3* and *cax1cax3* mutants, and CAX1 over-expressing Arabidopsis seeds (Punshon et al., 2012). Zn localization in the trichome cell itself is quite obvious in *A. halleri*, as the narrow Zn ring is localized more distal to the base, on the trichome stalk (Kupper et al., 2000;Zhao FJ, 2000;Sarret et al., 2009). In *A. lesbiacum*, Ni has a similar distribution along the stalk, and the authors suggested that the Ni could be stored inside vacuoles (Kupper et al., 2001). However, our mapping data from intact and sectioned trichomes of WT and OsZIP7-FOX leaves showed Zn in a ring shape at the base and continuing up into the trichome stalk (Figure 3), observations which are more consistent with an *extracellular* localization, because a vacuolar localization would presumably fill the trichome cell. In agreement with that, sectioned trichomes showed Zn at the edges of the cell (Figure 4), suggesting Zn localization in the apoplast (Gutierrez-Alcala et al., 2000). In *A. halleri* trichomes, Zn was previously shown to accumulate in a small compartment at its base. Whereas in roots of *A. halleri* Zn precipitates in the apoplast, in trichomes Zn might be in solid form as Zn oxides given the high concentration observed (> 1M when plants are treated with excess Zn), or in soluble form with O-donors such as citrate (Kupper et al., 2000), which could suggest extracellular localization (Sinclair and Kramer, 2012).

An alternative hypothesis is that Zn is localized in the cytoplasm, which would be compressed against the cell wall by the vacuole. In trichomes, cytoplasm stretches crossing the central vacuole are observed (Gutierrez-Alcala et al., 2000). Such stretches were observed in some sections, but did not correspond to Zn-rich regions (Figure 4D and 4E). Moreover, high Zn regions were also found at the interface of the trichome with mesophyll and socket cells, and at the apical side of socket cells to some extent (Figure 4D and 4E). If Zn was localized inside the tricome cell itself, we would expect a confined Zn distribution limited to the cytoplasm. Therefore, Zn is more likely to be in the apoplast of the trichome cells.

Previous work showed that Cd associated with trichomes was predominantly bound to oxygen (O) and nitrogen ligands in non-hyperaccumulators *A. thaliana* and *A. lyrata*, and in the hyperaccumulator *A. halleri* (Isaure MP, 2006;Fukuda N, 2008;Isaure et al., 2015). Cd may be associated with cell wall in trichomes, bound to O ligands (Isaure MP, 2006;Isaure et al., 2015). Similarly, in *A. halleri*, Zn was also shown to bind mainly to carboxyl and/or hydroxyl groups (Sarret et al., 2002). Phosphate, thiol and silanol groups were excluded as potential Zn ligands (Sarret et al., 2009). Interestingly, the non-hyperaccumulator *A. lyrata* showed more Zn bound to cell wall (40% of the Zn in trichomes) compared to hyperaccumulator *A. halleri* (20%) (Sarret et al., 2009). Thus, our data suggest that trichome-localized Zn is in the apoplasmic space, probably bound to the cell wall in *A. thaliana* trichomes.

Recently a new structure, the Ortmannian ring was described in *A. thaliana* trichomes (Kulich et al., 2015). The Ortmannian ring formation is dependent on the EXO70H4 exocyst subunit, and is a callose-rich secondary cell wall layer, localized between the basal and apical regions of the trichome stalk. Loss-of-function *exo70h4* plants showed no callose ring accumulation. Strikingly, WT and *exo70h4* differed in their ability to accumulate Cu in trichomes, with WT plants showing Cu accumulation at the base of the trichome, while *exo70h4* plants, which lack the Ortmannian ring, contain no Cu in the same region (Kulich et al., 2015). This data strongly suggest that the Ortmannian ring is involved in metal localization in trichomes. Therefore, Zn may be at the Ortmannian ring, or its distribution is being limited by it. Further investigations are needed to clarify this.

### Physiological significance of Zn accumulation in trichomes

We found that natural accessions of *A. thaliana* with higher whole leaf Zn have increased amounts of Zn in their trichomes (Figure 5 and Figure 6). When comparing percentage Zn in trichomes with whole leaf total Zn, we found a linear relationship: accessions with higher Zn concentration in whole leaves have increased Zn in trichomes (Figure 6). This leads us to hypothesize that trichome Zn accumulation might have a role in metal detoxification by providing a location for metal sequestration in *A. thaliana*.

Metal accumulation in trichomes has been shown before. In a previous study, four crop species where analyzed for Mn tolerance: sunflower (*Helianthus annuus*), white lupin (*Lupinus albus*), narrow-leafed lupin (*Lupin angustifolius*) and soybean (*Glycine max*). All but soybean could tolerate 100 µM Mn concentration without toxicity sympotoms (Blamey et al., 2015). Differently from the other Mn tolerant species, sunflower non-glandular trichomes accumulated Mn at the base, while Ca was distributed along the trichome length, resembling the pattern we observed in *A. thaliana* non-glandular trichomes with Zn and Ca. Fluorescence-XANES indicated that 66% of the Mn present at the base of the trichome is in the form of manganite (Mn(III)) (Blamey et al., 2015). Authors suggest that Mn is translocated from the apoplast to trichomes and then oxidized to manganite, avoiding toxicity to the cytoplasm and cell wall in leaf cells (Blamey et al., 2015). In the Ni-hyperaccumulator *Alyssum murale*, Mn excess was also shown to accumulate in trichomes, where it was associated with phosphorous (P) (MCNear, 2014). Thus, it is possible that Zn in *A. thaliana* trichomes is being transported to the trichome apoplast directly via the transpiration stream, as has been proposed for Mn (Blamey et al., 2015).

Hyperaccumulator *A. halleri* and non-hyperaccumulator *A. lyrata* also accumulate Zn at the base of trichomes (Kupper et al., 2000;Zhao FJ, 2000;Sarret et al., 2009;Isaure et al., 2015). It was also shown that Zn accumulates more in the mesophyll in *A. halleri* compared to veins, whereas the opposite is observed in *A. lyrata* (Sarret et al., 2009). In another study using XRF imaging of *A. halleri* leaves, Zn concentration in mesophyll cells increases 30-fold upon exposure of *A. halleri* plants to high Zn, whereas the trichome concentration only increases 3-fold (Kupper et al., 2000). These results indicate that, despite their high Zn accumulation, trichomes are not responsible for the Zn hyperaccumulation/hypertolerance traits in *A. halleri*. In tobacco (*Nicotiana tabacum*), the role of trichomes in heavy metal excretion is well documented, showing that both Zn and Cd are secreted from trichome tips as 20-150 µm crystals, where metals are present in Ca-containing compounds (Choi et al., 2001;Sarret et al., 2006;Isaure MP, 2010). However, these glandular, multicellular trichomes, are very different from the unicellular, non-glandular trichomes found in *A. thaliana* (Yang and Ye, 2013).

In *A. thaliana*, it has been shown that Cd and Mn accumulate at the base of trichomes (Ager FJ, 2003;Isaure MP, 2006). Interestingly, *A. thaliana* transgenic lines that accumulate varying concentrations of Cd showed trichome metal accumulation varying in a similar way: more Cd in leaves resulted in more Cd in trichomes (Ager FJ, 2003). This is consistent with what we observed for Zn in trichomes using 16 different accessions (Figure 5 and Figure 6). *A. halleri* trichomes accumulate Cd as shown by ^109^Cd autoradiography (Huguet S and I, 2012). Cd-enrichment in trichomes was clearly visible at three weeks of exposure time, but after nine weeks they could hardly be observed on the autoradiographs. These data indicate that Cd is first accumulated in trichomes, reaching a plateau, and then continues to increase in concentration in leaf tissues. As a consequence, trichomes could not be distinguished in autoradiographs from older leaves (Huguet et al., 2012). It is possible that Cd sequestration in trichome cells is a short-term response to metal excess, which becomes marginal upon continuous exposure, when leaf genes controlling hyperaccumulation/hypertolerance traits would be necessary. In agreement with that, *A. lyrata*, a non-hyperaccumulator, has more Zn in trichomes (20%) than in the hyperaccumulator *A. halleri* (10%) (Sarret et al., 2009). Thus, trichome metal sequestration and possibly detoxification might be more important in non-tolerant and non-accumulators *A. thaliana* and *A. lyrata* than in a metal hyperaccumulator, hypertolerant species.

Transgenic plants or natural accessions with increased Zn accumulation in leaves showed a higher percentage of Zn in trichomes. Therefore, our findings support that Zn found in trichomes increases linearly with higher Zn leaf concentrations (Figure 6). In *B. juncea*, experiments where mass flow is decreased by application of abscisic acid (ABA) to induce stomata closure showed that Cd accumulation in leaves is dependent upon transpiration, although root uptake is not affected (Salt et al., 1995). This would indicate that Cd accumulation in trichomes is dependent on transpiration rate. However, our data shows that Zn found in trichomes change depending on how much Zn a leaf accumulates, which suggest an active mechanism for Zn accumulation in trichomes. Whether this mechanism involves cell wall modifications or symplast transport remains to be answered.

## Conclusion

Our work provides evidence for Zn accumulation in trichomes of Arabidopsis thaliana, a genetically tractable model species, allowing exploration of the functional role of Zn distribution in trichomes and its relevance to leaf function. We also demonstrate that Zn accumulation in trichomes changes depending on Zn concentration in leaves, suggesting that plants might actively control this process to some extent. Future work should focus on the importance of such distribution in leaves and how it is molecularly controlled.

## Acknowledgements

This work was supported by NIH grant 5R01GM078536 and NSF Plant Genome grant DBI 0701119 to MLG and DES, and NIEHS grant ES007373 to MLG and TP. We also thank CNPq (Conselho Nacional de Desenvolvimento Científico e Tecnológico) for granting a fellowship to FKR.

This research used resources of the Advanced Photon Source, a U.S. Department of Energy (DOE) Office of Science User Facility operated for the DOE Office of Science by Argonne National Laboratory under Contract No. DE-AC02-06CH11357. Data was collected at APS beamline 2-ID-D. The assistance of Dr. Barry Lai and Dr. Stefan Vogt is gratefully acknowledged.

Use of the Stanford Synchrotron Radiation Lightsource, SLAC National Accelerator Laboratory, is supported by the U.S. Department of Energy, Office of Science, Office of Basic Energy Sciences under Contract No. DE-AC02-76SF00515. The SSRL Structural Molecular Biology Program is supported by the DOE Office of Biological and Environmental Research, and by the National Institutes of Health, National Institute of General Medical Sciences (P41GM103393). The contents of this publication are solely the responsibility of the authors and do not necessarily represent the official views of NIGMS or NIH. Data was collected at SSRL BL2-3. The assistance of Dr. Sam Webb and Dr. Ben Kocar is gratefully acknowledged.

## Notes

### Competing Interest Statement

The authors have declared no competing interest.

